# Mechanosensitive pore opening of a prokaryotic voltage-gated sodium channel

**DOI:** 10.1101/2022.05.10.491345

**Authors:** Peter R. Strege, Luke M. Cowan, Constanza Alcaino, Amelia Mazzone, Christopher A. Ahern, Lorin S. Milescu, Gianrico Farrugia, Arthur Beyder

**Author notes:** Corresponding authors: Gianrico Farrugia, M.D., Arthur Beyder, M.D., Ph.D. Mayo Clinic, 200 First Street SW Rochester, MN 55905., and; Lorin S. Milescu, Ph.D., University of Maryland at College Park College Park, MD 20742.

## Abstract

Voltage-gated ion channels orchestrate electrical activities that drive mechanical functions in contractile tissues such as the heart and gut. In turn, contractions change membrane tension and impact ion channels. Voltage-gated ion channels are mechanosensitive, but the mechanisms of mechanosensitivity remain poorly understood. Here, we leverage the relative simplicity of NaChBac, a prokaryotic sodium channel from *Bacillus halodurans*, to investigate its mechanosensitivity. In whole-cell experiments on heterologously transfected HEK293 cells, shear stress reversibly altered the kinetic properties of NaChBac and increased its maximum current, comparably to the mechanosensitive eukaryotic sodium channel Na_V_1.5. In single-channel experiments, patch suction reversibly increased the open probability of a NaChBac mutant with inactivation removed. A simple kinetic mechanism featuring a mechanosensitive pore opening transition explained the overall response to force, whereas an alternative model with mechanosensitive voltage sensor activation diverged from the data. Structural analysis of NaChBac identified a large displacement of the hinged intracellular gate, and mutagenesis at the hinge abolished NaChBac mechanosensitivity, further supporting the proposed mechanism. Overall, our results suggest that NaChBac responds to force because its pore is intrinsically mechanosensitive. This mechanism may apply to other voltage-gated ion channels, including Na_V_1.5.

## INTRODUCTION

Electrically excitable tissues with mechanical functions like the heart and gut use voltage-gated ion channels (VGICs) to generate electrical activity, which drives mechanical activity via electro-mechanical coupling^1^. Conversely, mechanical movements change membrane tension and impact electrical function in a process called mechano-electrical feedback^2^, which relies on specialized mechanically-gated ion channels, such as TREK^3^ and Piezo^4^. However, studies dating back nearly 40 years suggest that VGICs are also mechanosensitive and thus may directly contribute to mechano-electrical feedback^5–9^. Indeed, most VGIC families display mechanosensitivity, including sodium (Na_V_)^10^, potassium (K_V_)^11^, calcium (Ca_V_)^12^, proton (H_V_)^13^, and cyclic nucleotide-gated (HCN)^14^ channels.

Mechano-electrical feedback via VGICs can play a distinct physiological role. Unlike the specialized mechano-gated channels whose activation is generally voltage-insensitive, mechanosensitive VGICs create a “voltage-informed” mechano-electrical feedback^7, 15^. Perhaps the best example is the voltage-gated sodium channel Na_V_1.5, responsible for the upstroke of cardiac action potentials^16^. Given the heart’s role as a pump, Na_V_1.5 is a natural target for mechanosensitivity investigations, and several studies showed that macroscopic Na_V_1.5 currents are mechanosensitive^10, 17^. Interestingly, disease-associated Na_V_1.5 mutations (channelopathies) can affect mechanosensitivity^18–20^. Further, lipid-permeable anesthetics and amphipathic drugs such as ranolazine that target Na_V_1.5 inhibit its mechanosensitivity, often with little effect on its voltage-dependent gating^21, 22^. Despite this abundant phenomenological evidence, it is unclear whether mechanosensitivity is intrinsic to the channel or emerges through interactions with other factors, and the mechanism of mechanosensitivity in Na_V_ channels remains unknown.

Na_V_ channels operate through a complex gating mechanism, where the voltage-dependent movement of the four voltage sensors can trigger a voltage-independent physical opening of the intracellular gate in the pore, immediately followed by a fast and thorough inactivation^23^. Whether applied by fluid shear stress or membrane stretch, mechanical force alters the overall voltage sensitivity of macroscopic Na_V_ currents^8, 10, 17^, but we do not know how each gating transition is influenced by force. In principle, this information could be extracted by analyzing the response of single-channel events or macroscopic currents to mechanical stimuli, as recently shown for K_V_ channels^24^. However, the complexities of eukaryotic Na_V_ channel structure, together with its fast activation and inactivation kinetics, would make this mechanistic analysis challenging.

An alternative strategy is to use bacterial voltage-gated sodium channels, which have emerged as powerful models for eukaryotic Na_V_s^25^. Like their eukaryotic counterparts, prokaryotic Na_V_s are strongly voltage-sensitive^26^, have similar pharmacological sensitivities^27, 28^, and share some structural elements despite being homotetramers^25, 28, 29^. NaChBac from *Bacillus halodurans* is the first prokaryotic Na_V_ channel discovered^26^ and presents significant advantages for mechanistic studies: at one-fourth the coding sequence length of eukaryotic Na_V_s, NaChBac has simpler mutagenesis, structural symmetry, slower kinetics, and removable inactivation, which altogether facilitate detailed mechanistic investigations^27, 28^. In this study, we examined the mechanism of NaChBac mechanosensitivity through a combination of macroscopic and single-channel recordings, kinetic modeling, structural analysis, and mutagenesis and found that mechanosensitivity is intrinsic and likely resides with the channel pore.

## RESULTS

### Mechanical stimulation of bacterial voltage-gated sodium channels

We first tested if prokaryotic sodium channels are mechanically sensitive, as previously shown for eukaryotic Na_V_s^8, 10, 17^ (Figure 1). In a side-by-side comparison with the eukaryotic Na_V_1.5, we examined two prokaryotic channels: the wild-type (WT) NaChBac and a mutant (T220A) NaChBac with inactivation removed^27, 28^ (Figure 1A). We expressed each channel in HEK293 cells and assayed its mechanosensitivity via whole-cell electrophysiology, with fluid shear stress (1.1 dyn/cm^2^) applied as mechanical stimulation. Under control conditions, the wild-type NaChBac responded to depolarizing voltage pulses with steep activation followed by complete inactivation, like Na_V_1.5 but with slower kinetics (Figure 1B, Figure 1 Suppl A-D). The T220A mutant activated and stayed open with minimal inactivation (Figure 1B; Figure 1 Suppl. B).

**Figure 1.**
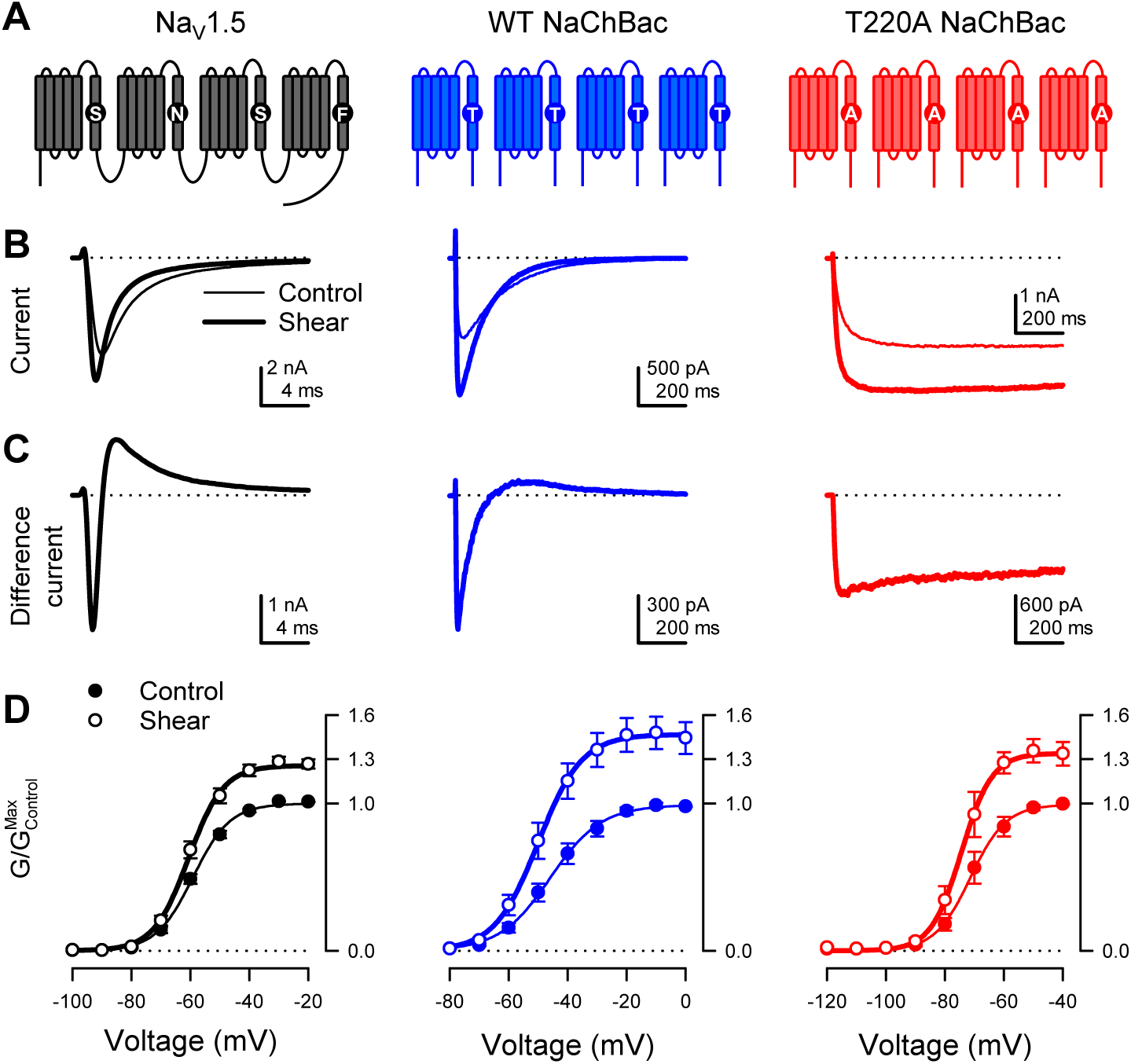
Shear stress increases the peak Na^+^ current of eukaryotic Na_V_1.5 and prokaryotic Na_V_ channel NaChBac. **(A)** Topologies of eukaryotic Na_V_ channel Na_V_1.5 (black) and prokaryotic Na_V_ channel NaChBac, without (WT, blue) or with (T220A, red) point mutation T220A, which makes NaChBac devoid of inactivation. **(B)** Representative Na^+^ currents elicited by a depolarization from -120 mV to -40 mV of Na_V_1.5 (black) and WT NaChBac (blue) or T220A NaChBac (red), before (—) or during (⁃) shear stress. **(C)** Difference currents obtained by subtracting the control trace from the shear trace in (B). **(D)** Voltage-dependent conductance normalized to the maximum conductance of controls (G/G_Max,Control_) for Na_V_1.5 (black), WT NaChBac (blue) or T220A NaChBac (red), before (—) or during (⁃) shear stress (n = 7-10 cells; P<0.05 by a paired 2-tailed t-test when comparing shear to control at voltages >-70 mV for Na_V_1.5, >-60 mV for WT and >-80 mV for T220A).

Shear stress increased the whole-cell currents of both prokaryotic channels, comparably to Na_V_1.5 (Figure 1B, “control” vs. “shear”; Figure 1 Suppl. B, E; I_Peak_ in Table 1). Both activation and inactivation responded to shear stress, as demonstrated by the difference currents (I_Shear_ – I_Control_) from both wild-type NaChBac and Na_V_1.5 (Figure 1C). Removal of inactivation in NaChBac T220A allowed us to separate these responses and focus on activation. Shear forces also increased T220A NaChBac currents, albeit slightly less than wild-type (Figure 1C), suggesting that mechanical forces act predominantly on the mechanistic steps associated with the channel’s activation and/or opening. Overall, shear stress increased maximum conductance (G_Max_) by 47% for WT NaChBac and 34% for T220A NaChBac, compared to 26% for Na_V_1.5 (Figure 1D, G_Max_ in Table 1).

**Table 1.**
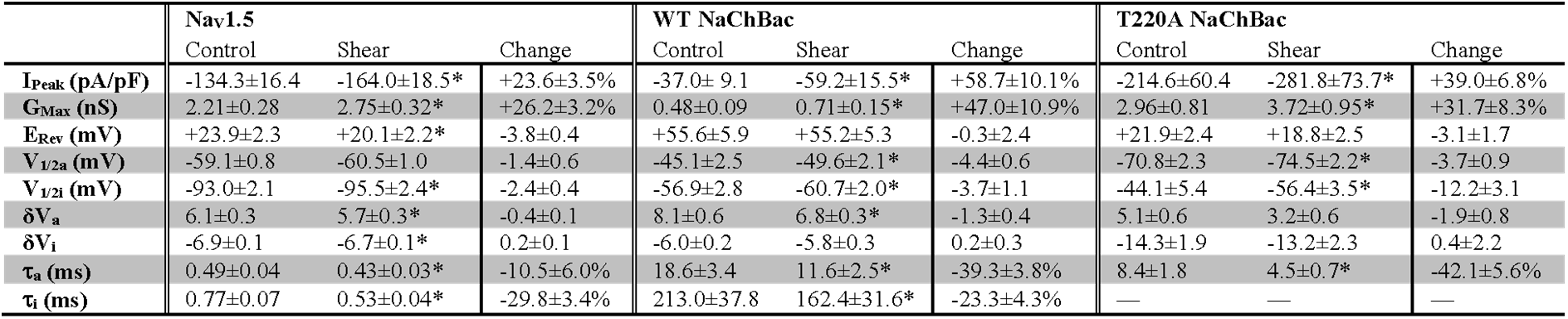
Effect of shear stress on parameters of wild-type and T220A NaChBac. Shear, flow of extracellular solution; I_Peak_, maximum peak current density; G_Max_, maximum peak conductance; E_Rev_, reversal potential; V_1/2a_, voltage dependence of activation; δV_a,_ slope of steady-state activation; V_1/2i_, voltage dependence of inactivation; δV_i,_ slope of steady-state inactivation; τ_a_, time constant of activation at -30 mV; τ_i_, time constant of inactivation at -30 mV. The background of Na_V_1.5 was H558/Q1077del. Number of cells: Na_V_1.5, 10; wild-type (WT) NaChBac, 7; T220A NaChBac, 7; T220A/I228G NaChBac, 11. \**P*<0.05 shear *vs.* control by a two-tailed paired Student’s t-test.

Although the steady-state conductance curves obtained under shear stress mostly appear as vertically stretched versions of the control curves, accounting for the higher maximum current, they exhibit a slight negative shift of the half-activation voltage (Figure 1D; V_1/2a_ in Table 1). This effect is more easily visualized when each conductance curve we normalize to its maximum (Figure 1 Suppl. F). Shear stress also increased the conductance slope (δV_a_ in Table 1), while the foot of the activation curve did not change. Interestingly, the inactivation voltage mid-point also shifts negative (Figure 1 Suppl. G; V_1/2i_ in Table 1). Kinetically, shear stress accelerates the time course of both activation (Figure 1 Suppl. C; τ_a_ in Table 1) and inactivation (Figure 1 Suppl. D; τ_i_ in Table 1).

### Interactions between electrical and mechanical stimuli

The whole-cell shear stress experiments demonstrate that mechanical forces affect NaChBac macroscopic currents, and these results are likely to have mechanistic implications. However, ambiguities inherent to macroscopic currents limit the information that can be extracted from data about individual state transitions. Hence, we addressed these ambiguities via single-channel recordings before conducting a mechanistic analysis to determine how force interacts with voltage to gate the channel. To simplify experiments and interpretations, we focused on NaChBac T220A, which lacks inactivation^27, 28^. We expressed NaChBac T220A in Piezo1-knockout (P1KO) HEK293 cells, free of mechanosensitive channel activity^30^ (Figure 2A, Fig 2. Suppl. A-F). We assayed mechanosensitivity via cell-attached patch-clamp electrophysiology, using a high-speed pressure- clamp^31^ to apply controlled suction to patches.

**Figure 2.**
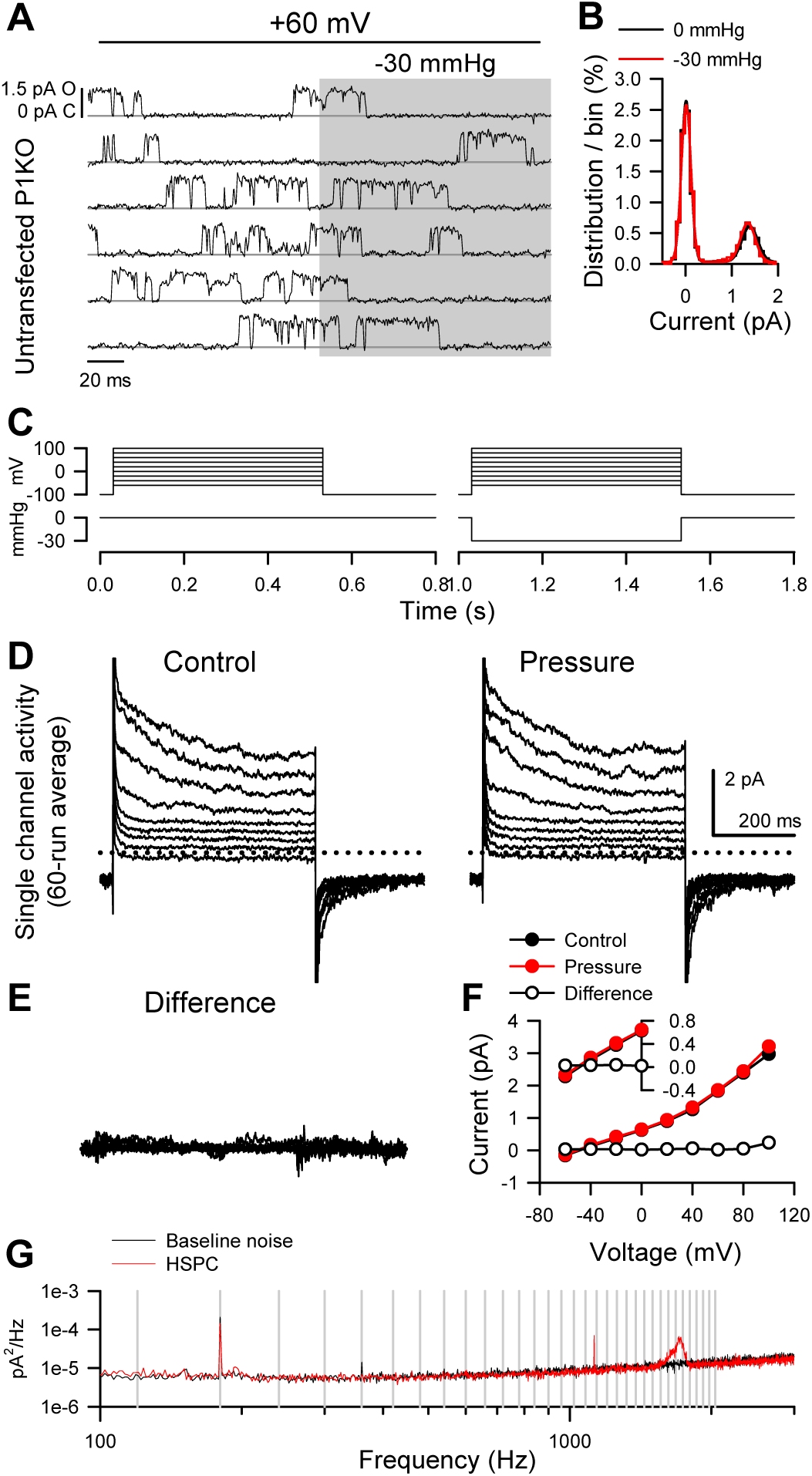
Patch pressure increases the open channel probability of T220A NaChBac single channels in P1KO cells. **(A)** Representative traces of single T220A NaChBac channels at -80, -60, -40, or -20 mV and with 0 (unshaded) or -10 mmHg (shaded region) applied to the patch. **(B)** All-point histograms constructed from the traces shown in (a) at -80, -60, or -20 mV and 0 (black) or -10 mmHg (red) binned every 0.2 pA. Bins were normalized to an area of 1 and fit with a sum of two Gaussians, in which open events at -60 mV were 0.77 pA and 0.17 P_O_ without pressure and 0.75 pA and 0.72 P_O_ (330% increase) with pressure; open events at -20 mV were 0.43 pA and 0.90 P_O_ without pressure and 0.42 pA and 0.90 P_O_ (0% increase) with pressure. **(C)** Mean open probabilities (P_O_) at voltage steps from -100 to -20 mV with 0 (black) or -10 to -50 mmHg (red gradient) pressure (n=7-21 cells per voltage; *P<0.05, control vs. pressure by a paired 2-tailed t-test). **(D)** P_O_ per voltage from (C), re-plotted versus pressure (0 to -50 mmHg).

The single-channel amplitude of voltage-gated sodium channels is tiny (∼1 pA at -80 mV and ∼0.5 pA at -20 mV), and pressure-clamping introduces additional noise and transient artifacts. Together with rapid kinetics, these limitations have traditionally prevented single-channel studies on mechanosensitivity in VGICs. After careful mechanical and electrical optimization, despite the low signal-to-noise ratio typical for sodium channels^32^, and the noise introduced by the pressure clamp (Figure 2 Suppl. G), we were able to resolve single-channel events across a physiologically relevant voltage range, and with enough bandwidth (∼1 kHz) to capture sufficiently fast kinetics (Figure 2A).

Suction on the membrane patch exerts a mechanical force on the channel^33^. Because patches have non-zero resting tension^34^, we designed stimulation protocols to test voltage- and mechano-sensitivity in a pairwise fashion (Figure 2A), enabling us to compare suction-induced changes to a no-suction baseline for all channels and traces. Within each 400 ms voltage step from -100 to -20 mV, the suction pressure alternated between 0 and -10, -30, or -50 mmHg. Thus, we could obtain and compare control and pressure data in the same cell, using test pressures relevant for mechanosensitive channel function^33, 35^. As indicated by the current amplitude histograms (Figure 2B), the single-channel current is less than 0.5 pA at -20 mV, but we could still separate the closed and open levels. Above -20 mV, the unitary current became too small for reliable analysis. Using a half-amplitude threshold method, we measured open state occupancy between -100 and -20 mV (Figure 2C). We cross-checked this approach against fitting all-point amplitude histograms with sums of two Gaussian distributions, one for each current level (Figure 2B), where the relative weight of the open-level Gaussian indicates the open state occupancy probability (P_O_). The two methods produced similar results.

Under control conditions (zero applied patch pressure), P_O_ was strongly voltage-dependent (Figure 2A-C), as predicted by the whole-cell activation curve (Figure 1D). P_O_ was nominally zero at -80 mV and below, and P_O_ increased as the voltage became more positive, reaching 0.525 at -20 mV. Relative to whole-cell activation, the P_O_ curve is shallower and ∼20 mV more positive. This discrepancy is likely an artifact of a variable and non-zero resting potential, unmeasurable in cell-attached recordings (averaging sigmoid curves with a scattered and shifted midpoint results in a shallower and shifted sigmoid).

Patch suction altered the voltage-dependent P_O_ (Figure 2A-C; Table 2). At extremely negative voltages (-100 and -80 mV), where the channel is closed under control conditions, P_O_ remained zero with suction. However, pressure significantly increased P_O_ at more positive voltages. Responses were dependent on suction strength (Figure 2C, D), but even at high negative pressures (-30 and -50 mmHg), the induced changes were confined to the voltage activation range (-80 to -20 mV) (Figure 2C, D). These results agree with the whole-cell experiments, where shear stress did not move the foot of the activation curve but stretched the curve vertically. As single-channel data yield the actual P_O_ values under different pressures and voltages, we could establish that the increase in whole-cell conductance results from an increase in P_O_ and not in single-channel conductance, which remained constant under pressure (Figure 2A, B).

**Table 2.**
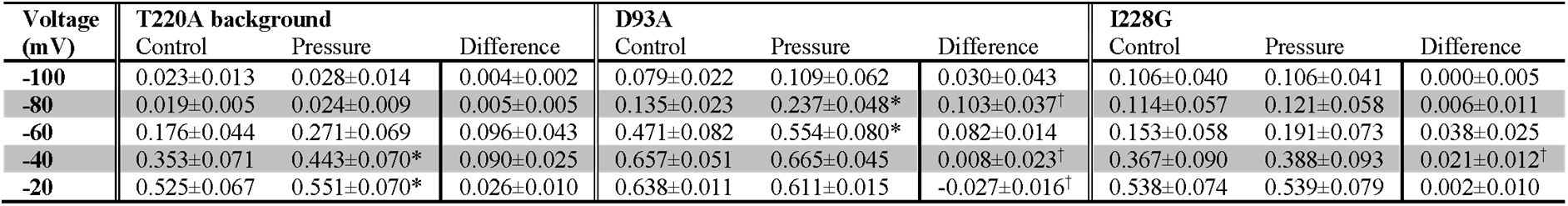
Effect of pressure on open probability of mutants D93A and I228G in the T220A NaChBac background. Open probability; *n* = 6-12 cells; *P<0.05, -10 *vs.* 0 mmHg pressure, by a 2-tailed paired t-test; ^†^P<0.05, D93A or I228G *vs.* T220A background by a 2-tailed unpaired t-test.

Because some previous studies have shown that shear stress and patch pressure can create irreversible changes^11, 17, 36^, we tested specifically for reversibility in our preparations. In whole-cell experiments, we found that the increase in peak NaChBac T220A current density induced by shear stress is fully reversible (Figure 3A-B). With single channels, to test the reversibility of P_O_ increase by patch pressure, we lengthened the time before pressure application to 2 s, applied -30 mmHg pressure for 500 ms, and compared the pre- and post-pressure Po values (Figure 3C, Figure 3 Suppl. A). Pressure increased P_O_ throughout the -80 to -20 mV activation range (Figure 3 Suppl. B), with 20 out of 21 cells responding at -60 mV (Figure 3D-E). Once pressure returned to 0 mmHg, P_O_ returned to its baseline value (Figure 3F, Figure 3 Suppl. C-D). As expected, this change was not instantaneous because the channel must transition back into a different set of state occupancies, which takes time (Figure 3 Suppl. B).

**Figure 3.**
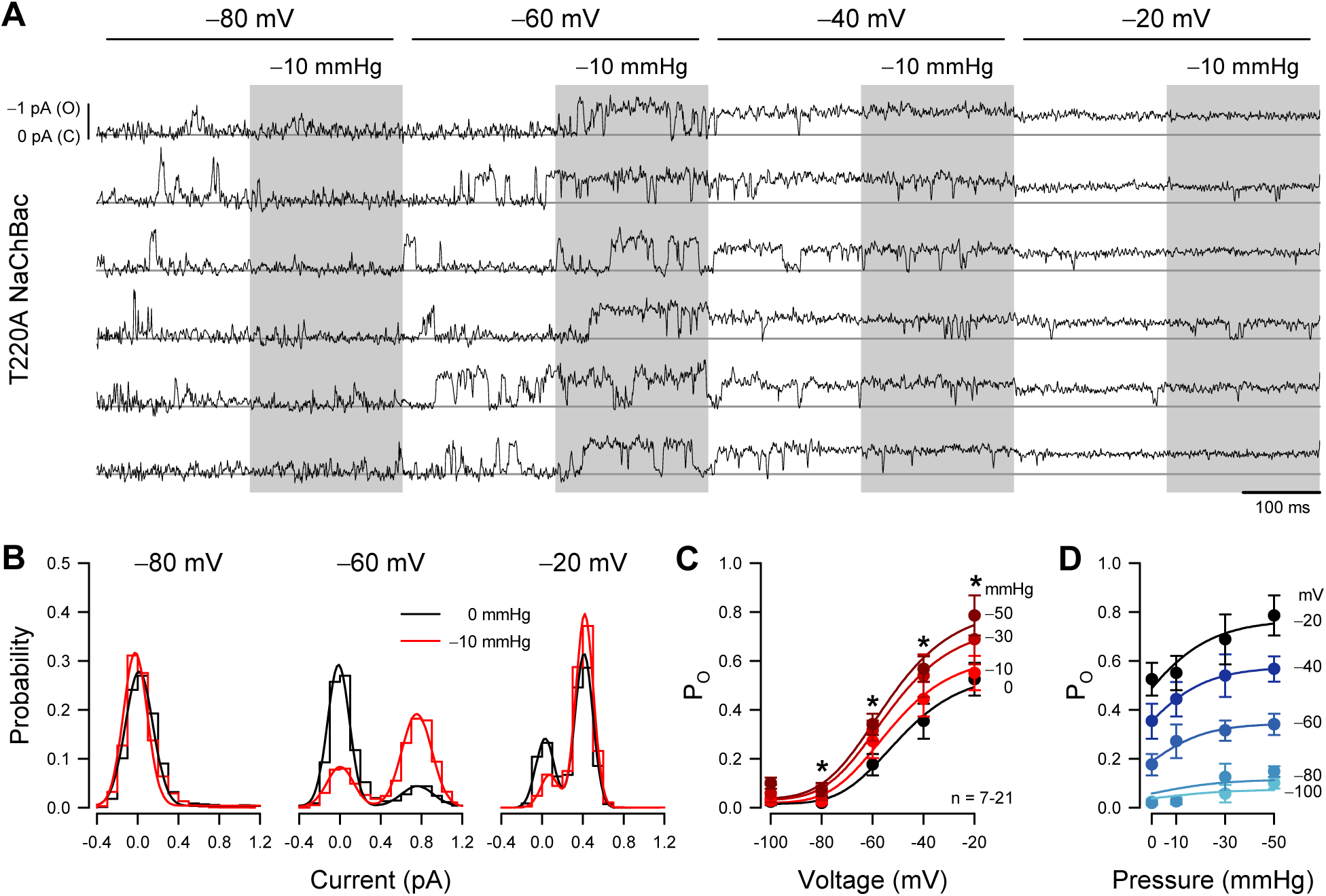
Pressure-sensitive increase in whole-cell peak currents and single-channel open probability of T220A NaChBac is reversible. **(A)** Representative whole cell currents from HEK cells expressing T220A NaChBac were elicited by a voltage protocol (Figure 1 Suppl. A) before (black), during (red), or after (blue) shear stress. **(B)** Peak current densities before (black), during (red), or after (blue) shear stress (n = 5 cells, *P<0.05 to pre-control by a one-way ANOVA with Dunnett’s post-test). **(C)** Representative single channel activity at -60 mV from Piezo1-knockout HEK cells transfected with T220A NaChBac, before (unshaded), during (shaded region), or after application of -30 mmHg to the patch for 500 ms. **(D)** All-sample distributions of single channel activity from the cell shown in (C), binned every 0.05 pA with peaks at 0 pA (closed) and ∼0.9 pA (open). **(E)** Mean open channel probability (P_O_) per cell (gray circles) before (black), during (red), or after (blue) application of -30 mmHg pressure. **(F)** Differences in post-pressure P_O_ (ΔP_O_) from pre-pressure controls.

### Mechanical force mainly affects pore opening

An obvious interpretation of the whole-cell and single-channel results is that force alone does not open the channel. If it did, we would see openings at voltages where the channel is typically closed, provided that we applied enough membrane tension. Instead, we see that force enhances openings (increases P_O_) that are already driven by membrane depolarization. A simple interpretation is that force does not create additional conformational states but modifies the energetics of the existing transitions. If this is true, then force will interact with at least one mechanistic component: (1) voltage sensor activation, (2) pore opening, or (3) inactivation. We consider that inactivation is unlikely to play a significant role. First, NaChBac T220A responds to patch pressure like the wild type, even though the mutant virtually lacks inactivation (Figure 1B, C). Second, eukaryotic Na_V_ and wild-type NaChBac have similar responses to shear stress (Figure 1B and C), even though they inactivate via different mechanisms^37^. Thus, the effects of force on inactivation could simply be due to the coupling of inactivation to activation^38^. For these reasons, we focus here on the NaChBac T220 channels, which show minimal inactivation.

The remaining possibilities are that force interacts with (1) voltage sensors or (2) the pore. While not necessarily mutually exclusive, the two extreme models corresponding to these interactions are easier to formulate and discriminate than mixed models. Hence, we examined specific changes in kinetic properties from the force and compared them against model predictions. We first formulated a kinetic model (Figure 4A) that encapsulates the homo-tetrameric nature of NaChBac T220A, its voltage-dependent activation, and its lack of inactivation. We made the rates along the activation pathway (closed states C_1_ to C_5_) strongly voltage-dependent to agree with the whole-cell and single-channel activation curves (Figures 1D and 2C). In contrast, we made the concerted opening transition (C_5_ to open state O_6_) voltage*-*independent, as previously shown^39^ and based on our observation that the whole-cell activation curve reaches a steady maximum (Figure 1D), which, according to the single-channel data, corresponds to a maximum P_O_ of ∼0.6 (Figure 2C). If the concerted opening were significantly voltage-dependent, the maximum P_O_ would approach unity at strongly depolarizing voltages. The model parameters were manually optimized to match the experimental data under control conditions (see Methods).

**Figure 4.**
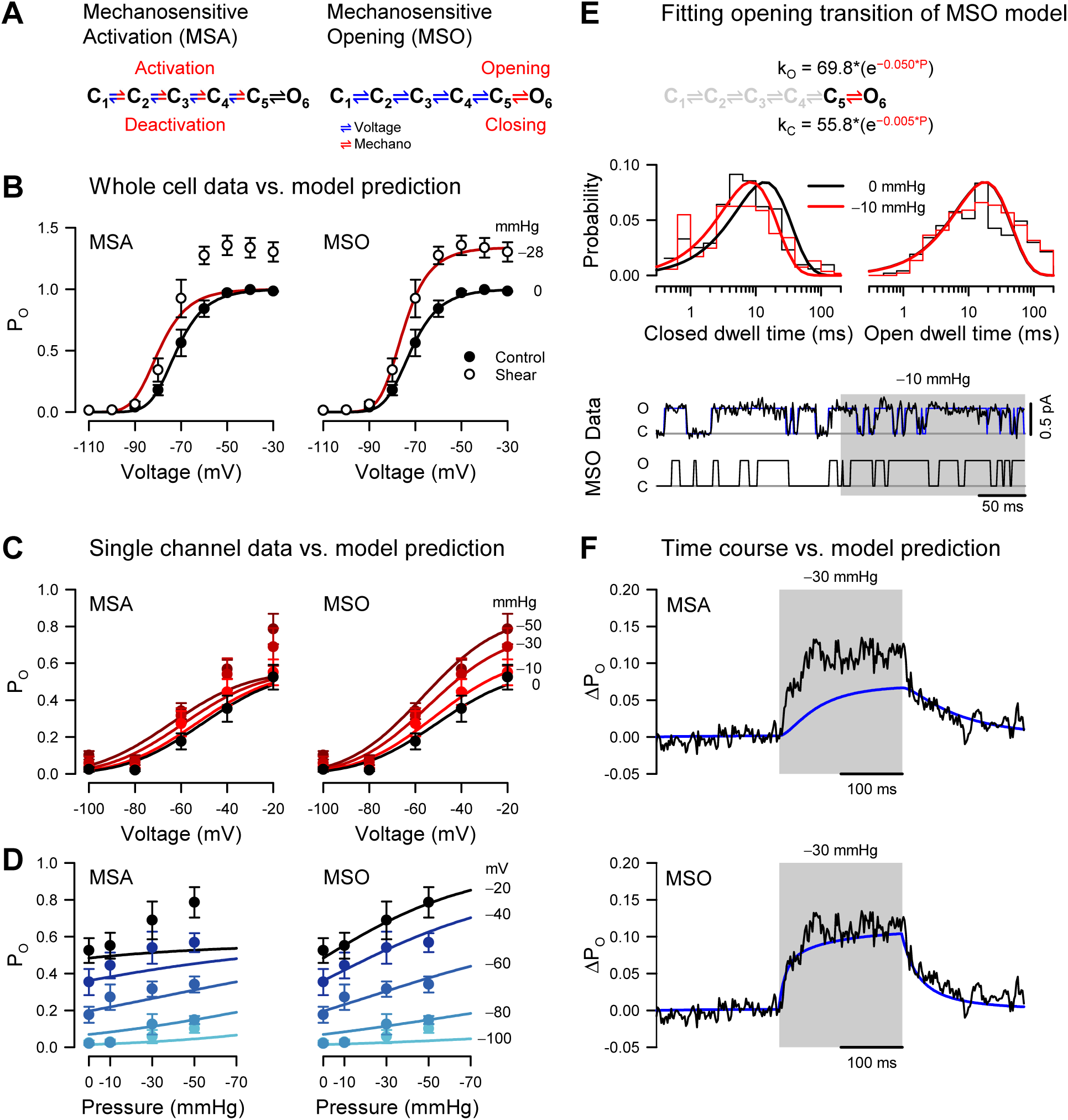
Pressure destabilizes the T220A NaChBac closed state. **(A)** Mechanosensitive activation (MSA) depicts a model in which the C_1_ to C_5_ closed state transitions are both voltage- and pressure-dependent (blue and red); mechanosensitive opening (MSO) depicts a model in which the C_1_ to C_5_ closed state transitions are voltage-dependent (blue), and the C_5_ closed to O_6_ open state transition is pressure-dependent (red). Rate constants: *k_a_* = 800 × *e*^0.0055×*V*^, *k_d_* = 0.1 × *e*^0.0055×*V*^. **(B)** MSA (left) and MSO (right) model predictions of open probability (P_O_) across voltages from -110 to -30 mV with 0 (black) or -28 mmHg applied pressure (dark red), compared to G/G_Max_ whole cell data (Figure 1D) with 0 (●) or 10 mL/min (○) fluid shear stress. **(C-D)** MSA (left) and MSO (right) model predictions of single channel P_O_ (●) plotted versus voltage (C) at pressures from 0 to -50 mmHg (red gradient) or versus pressure (D) at voltages from -100 to -20 mV (blue gradient). **(E)** MSO model adapted fit to a single pressure-sensitive C_5_ to O_6_ transition with pressure-dependent kinetic constants assigned for opening (*k*_O_) and closing (*k*_C_). Insets: top, closed (left) and open (right) dwell time histograms of single channel data versus the MSO model PDF curves; bottom, single channel trace recorded at -20 mV (black) and idealization (blue) with -10 mmHg applied to the region shaded (gray), compared to a trace from the MSO model. **(F)** MSA (top) and MSO (bottom) model prediction (blue) of single channel P_O_ at -60 mV before, during, and after pressure, compared to the average current from single channel data (black).

### Mechanosensitive activation

The first scenario, where mechanical force interacts only with the voltage sensors, is captured by a mechanosensitive activation (MSA) model (Figure 4A). In this case, we expect to see force-induced changes in the mechanosensitive rate constants along the C_1_ to C_5_ pathway. Experimentally, we observed increased whole-cell current by shear stress (Figure 4B), matched by an increase in P_O_ when membrane tension is raised via patch suction (Figure 4C). With the MSA model, we can explain this result by ascribing positive tension sensitivity (i.e., negative pressure sensitivity) to the activation (forward) rates and/or negative tension sensitivity to the deactivation (backward) rates. A situation where both activation and deactivation rates have positive or negative tension sensitivities is acceptable, as long as the forward sensitivities are more positive than the backward ones.

The MSA model predicts that the activation curve shifts toward more negative voltages when tension increases, but its slope and maximum value remain precisely the same (Figure 4B, MSA). The activation midpoint would change because tension shifts the equilibrium of each activation step (C_1_ to C_5_) toward C_5_ at any given voltage. In contrast, the slope and maximum P_O_ would be unchanged by tension because they are determined by the voltage-sensitivity of activation and by the voltage- and force-independent opening transition (C_5_ to O_6_), respectively. In other words, extreme tension would push the channel to reside in the C_5_ and O_6_ states, but the equilibrium between these two states – and hence maximum P_O_ – would remain the same. However, we did not observe this behavior experimentally. Instead, when membrane tension increased, both the whole-cell activation curve (Figure 4B) and the P_O_ curve (Figure 4C) exhibited increased steepness and greater maximum value, while the foot of each curve remained approximately unchanged. The experimental activation data are thus in stark contrast with the predictions of the MSA model.

### Mechanosensitive opening

The alternative scenario, where mechanical force interacts only with the channel pore, is captured by a mechanosensitive opening (MSO) model (Figure 4A). In this case, we expect to see force-induced changes in the mechanosensitive C_5_ to O_6_ rate constants. With the MSO model, the observed increase in P_O_ by tension can be explained by ascribing positive tension sensitivity to the opening (forward) rate, and/or negative tension sensitivity to the closing (backward) rate, or any combination where the forward sensitivity is more positive than the backward one.

The MSO model predicts that the activation curve reaches a larger value and becomes steeper when tension increases and shifts slightly toward more negative voltages, with a relatively unchanged foot (Figure 4B, MSO). The maximum P_O_ would change because it is determined by the tension-dependent pore opening and closing rates, but why would the voltage activation curve shift and steepen under tension when the tension-dependent rates are voltage-insensitive? To understand this, we must consider the final two states together, C_5_ and O_6_. Their joint occupancy is determined by the voltage-dependent but tension-independent activation/deactivation rates. Thus, membrane tension would not alter the voltage-dependent profile of the joint C_5_ and O_6_ occupancy but would change the occupancy ratio between these two states, favoring the open state. Therefore, the greater the joint occupancy, the greater the P_O_ at any given voltage. The result would be an asymmetrical shift in the activation curve at the top versus the bottom, increased steepness, and a greater maximum value. Indeed, the MSO model supports the mechanically induced changes in the whole-cell and single-channel activation curves (Figure 4B and C, MSO).

Having examined the changes in P_O_ vs. voltage under different tension values, we conversely examined P_O_ vs. tension under different voltages (Figure 4D). Reversing voltage and tension as independent variables does not create new information, as we are using the same data points as in Figure 4C, but it makes it easier to judge the fitness of each model. Thus, the MSA model predicts a significant shift in the P_O_ vs. tension curve when the voltage increases but no change in the maximum value and the slope of the curve (Figure 4D, MSA). In contrast, the MSO model predicts a significant change in the maximum value and the slope but only a small shift in the curve and a slight change in its foot (Figure 4D, MSO). The experimental P_O_ data points align well with either the MSA or the MSO model at zero pressure. However, the MSO model becomes a significantly better match to the data as the pressures increase (Figure 4C).

### Mechanical force destabilizes the NaChBac closed state

The analysis so far clearly favors the MSO model. However, we used only the steady-state information in the data, and we do not know if the MSO model can also explain the observed kinetics. The MSO model assumes tension-dependent opening and closing rates (at least one, if not both), whereas the MSA model assumes these rates to be tension-independent. If the pore opening transition were tension-dependent, then the pore opening (C_5_ to O_6_) and/or the closing (O_6_ to C_5_) rate would be affected by force, which would be reflected in the single-channel closed and open lifetimes. In our simple NaChBac kinetic model, the open state lifetime distribution has only one component, with the time constant equal to the inverse of the closing rate constant (O_6_ to C_5_). In contrast, the closed state lifetime distribution has five components, without an easy way to isolate the opening rate constant. However, the deactivation rates are likely so small at extremely depolarizing voltages (e.g., ≥-20 mV) that the channel essentially flickers between the last two states (C_5_ and O_6_). Hence, as an approximation, the closed lifetime distribution has one component, and its time constant approaches the inverse of the opening rate constant (C_5_ to O_6_). Consequently, a truncated model with only the final two states would approximate the channel at -20 mV (Figure 4E).

Because NaChBac T220A has some residual inactivation (Figure 1 Suppl. E, J), we used relatively short (200-500 ms) voltage/pressure stimulation episodes, so many recorded traces contained no events. To fit the single-channel data with the MIL algorithm^40^, we had to discard the first and last dwells in each trace because they are likely truncated and cannot be used for analysis, which means that all the event-less traces were also discarded. Under these conditions, the remaining data would yield a significantly higher P_O_ and bias the estimated rates. To partially compensate, we constrained the model parameters^41, 42^ to enforce a ratio between the opening and closing rate constants corresponding to the P_O_ measured under control (zero added tension) conditions. We also constrained the pressure sensitivity parameters since we can reliably estimate them from the P_O_ data, but we verified that we could obtain similar results without this constraint. Although the single-channel fits are subject to inherent stochasticity (Figure 4E), they clearly show that, under tension, the closed state lifetime distribution shifts toward shorter dwell times. The average closed lifetime approaches the bandwidth limit (∼1 ms), but the fitting algorithm partially compensates for the missed events. In contrast, the open state distribution remained virtually unchanged by tension.

The observed shift in the closed state lifetimes further confirms that the channel is better represented by the MSO model, as the competing MSA model would exhibit no such shift at saturating voltages. Moreover, it suggests that force destabilizes the closed state as the opening rate changes with tension. As we now have an idea about the magnitude of opening and closing rates, we can also examine activation kinetics. In principle, we can extract this information by fitting the single-channel data recorded at intermediate voltages (e.g., -60 mV), where the channel visits all states. However, the changes in voltage and pressure stimuli make these data non-stationary, and a more straightforward approach is to examine the macroscopic data created by averaging the single-channel recordings. As shown in Figure 4F, the MSO model captures well the time course of the average current and gives us an idea about the magnitude of the activation rates. In all, our modeling of the whole cell and single channel results suggest that the MSO model, which assigns tension sensitivity to the voltage-insensitive pore opening step, best fits the experimental data and places the NaChBac mechanosensor within the pore.

### Pressure affects the stability of the intracellular gate

According to the “force-from-lipid” model^43^, ion channels gain mechanosensitivity when their cross-section expands or shrinks upon a conformational change^44, 45^. Based on our kinetic analysis, the site of mechanosensitivity in NaChBac is most likely the pore opening, the final gating transition (C_5_ to O_6_ in the MSO model in Figure 4A). Interestingly, previous structural modeling studies have predicted that when voltage sensors are suitably activated, mechanical energy is required to open the gate^46^, which implies that negative membrane tension (i.e., patch suction) would facilitate opening. If our hypothesis were true, we would predict a change in the cross-section between the final two states in the MSO model: the activated but still closed C_5_ and the open O_6_. To test this hypothesis, we examined the two existing prokaryotic voltage-gated sodium channel structural models: Na_V_Ab, capturing the channel in the closed conformation^47^, and Na_V_Ms, representing the open state^48^.

By contrasting closed and open models, we searched for the channel substructures undergoing the largest movements within the membrane plane and found that the intracellular portion of the pore-forming S6 segment is displaced laterally around a “gating hinge” (Figure 5A, B). Interestingly, this type of movement has been previously proposed in functional studies^49–51^ and confirmed by structural experiments,^52^ including an example where the intracellular side of a voltage-gated ion channel pore was found to expand the area of the bilayer’s inner leaflet upon S6 lateral movement^49, 53^.

**Figure 5.**
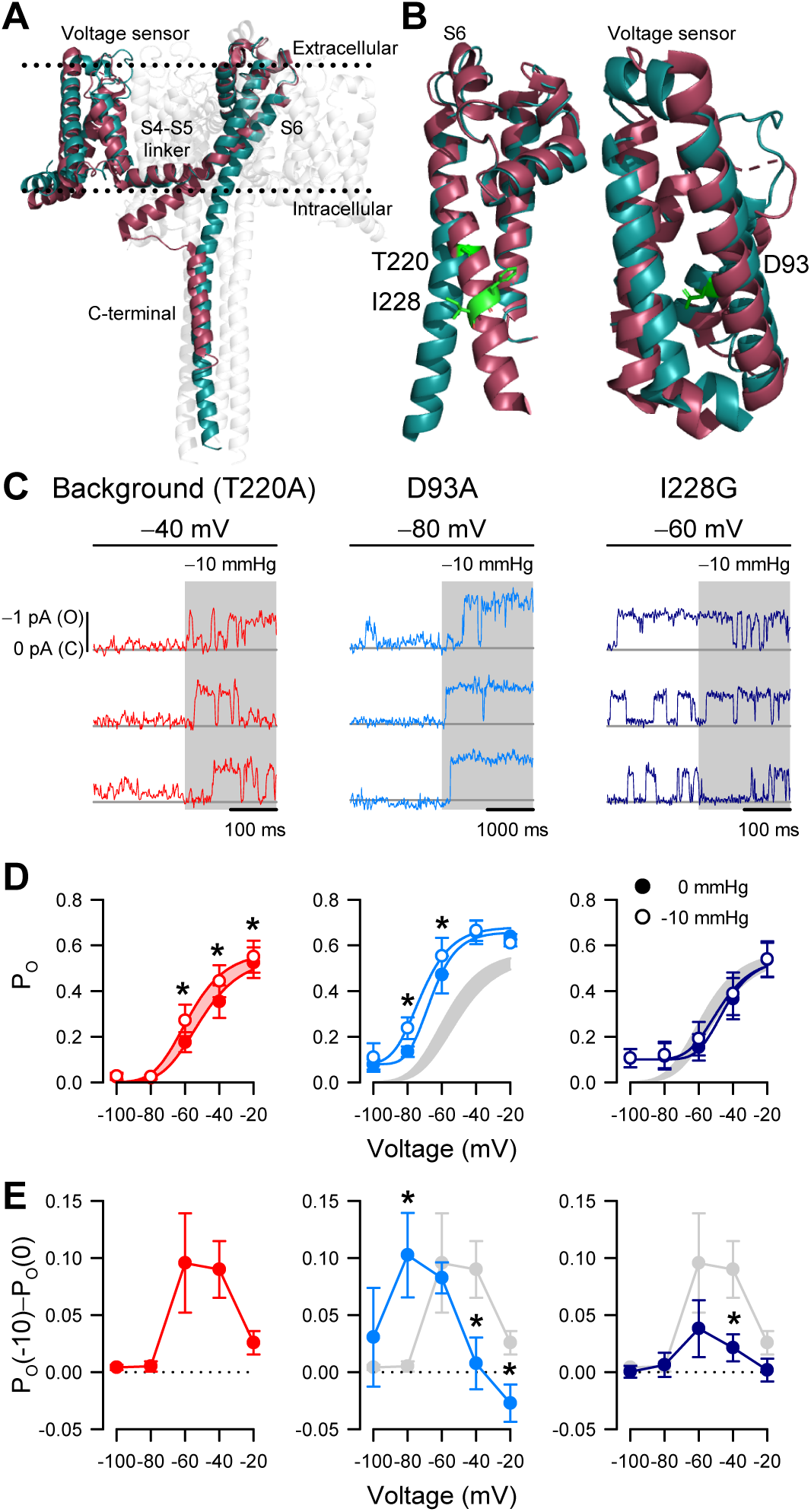
I228G disrupts the pressure sensitivity of NaChBac background T220A. **(A)** Conformational change of prokaryotic Na^+^ channels from the closed (cyan, Na_V_Ab, 2017) to open state (magenta, Na_V_Ms, 2017), illustrating the movement of the voltage sensor, S4-S5 linker, S6 segment, and C-terminal tail in relation to the lipid bilayer. **(B)** Location of key residues T220A and I228 in the S6 pore segment and D93 in the voltage sensor. **(C-D)** Voltage-dependent open probabilities ((D), P_O_) of single channel activities (C) recorded at the indicated voltages with 0 or -10 mmHg pressure from P1KO cells expressing the T220A NaChBac background (red or gray shading) or with additional mutations D93A (blue) or I228G (indigo). (*P<0.05, -10 mmHg vs 0 mmHg, n = 338-636 traces per voltage from 6-12 cells). Half-points of open probability (0 *vs.* -10 mmHg): T220A, -45.6 *vs.* -58.1 mV; D93A, -65.1 *vs.* -72.3 mV; I228G, -42.1 *vs.* -43.9 mV. **(E)** Difference in open probability induced by -10 mmHg pressure (P_O_(-10)–P_O_(0)) as a function of voltage in the control background (red or gray shading) or with D93A (blue) or I228G (indigo).

According to our structural analysis, the “force-from-lipid” model applies to NaChBac. Because S6 helices move and open the pore only after voltage sensors activate, it follows that mechanosensitivity, which is associated with S6 movement, resides with the pore opening (C_5_ to O_6_ in the MSO model) and not with the voltage sensor activation (C_1_ to C_5_ in the MSA model). This interpretation agrees with the MSO model, with one potential caveat being that Na_V_Ab as a closed channel could represent other closed states along the activation pathway, rather than the fully activated closed conformation (C_5_ in the MSO model). As a result, a mechanosensitive transition could still occur before pore opening. In other words, although the pore opening is likely mechanosensitive, it might not be the only mechanosensitive transition based solely on these structural models.

If mechanosensitivity were exclusive to pore opening, preventing S6 lateral movement via mutagenesis would abolish the effects of patch suction on P_O_. However, if voltage sensor activation were also mechanosensitive, then voltage sensor mutagenesis would only change the response to suction but not eliminate it. We tested these ideas via site-directed mutagenesis within the S6 hinge and the voltage sensor, using NaChBac T220A as background. Most mutations we tried within the pore resulted in non-expressing or non-functional channels, but eventually, we identified I228G in the S6 hinge region (Figure 5B). Within the voltage sensor, we chose D93A to stabilize the sensor in the resting position^54^ (Figure 5B). We applied the same single-channel experimental paradigms to directly compare the double mutants (NaChBac T220A plus I228G or D93A) with the T220A results described above (Figure 5C).

The voltage sensor NaChBac T220A+D93A double mutant shifted its voltage sensitivity relative to T220A (Figure 5D; Figure 5 Suppl. C). However, its mechanosensitivity remained intact and followed the negative shift of voltage-dependent gating (Figure 5D, E). The pore NaChBac T220A+I228G double mutant channel exhibits some interesting properties. First, the channel could gate normally with voltage, like the single mutant controls (Figure 5D). However, P_O_ did not reach zero at -80 mV but remained around 0.1. Second, the effect of membrane tension on P_O_ was nearly eliminated (Figure 5E). Thus, at -60 mV, membrane tension increased P_O_ by 0.096 for the NaChBac T220A mutant but only by 0.035 for NaChBac T220A+I228G, corresponding to ∼3- fold difference in effects between the two mutants. At -40 mV, the difference was even more significant (∼4-fold): 0.090 with NaChBac T220A and only 0.025 with NaChBac T220A+I228G. We could explain the small remaining effect of tension on P_O_ in the double mutant in two ways: either cross-section expansion due to a partial displacement of S6 during pore opening or another weakly mechanosensitive transition in the gating mechanism. The first possibility seems more plausible because some degree of S6 displacement is probably necessary for channel opening, and also because NaChBac T220A+D93A maintained a tension sensitivity similar to NaChBac T220A, even though its voltage sensitivity shifted by more than -30 mV (Figure 5D). Overall, these mutagenesis results provide experimental evidence that strengthens our conclusion that mechanical forces interact primarily with the pore opening transition.

## DISCUSSION

Electrically excitable cells depend on concerted efforts by voltage-gated ion channels (VGICs) to detect small changes in transmembrane voltage and amplify them to produce a wide range of action potentials^55^. Some electrical organs, such as the heart, bladder, and gut, function primarily as mechanical pumps, using excitation-contraction coupling to drive muscle contractions. Cells in these pumps experience significant recurrent changes in membrane tension that can potentially affect the activity of membrane proteins, which, in turn, can affect organ function by a process called mechano-electrical feedback^7, 8, 15, 56^. For VGICs in mechanical environments, mechanosensitivity may integrate both electrical^57^ and mechanical signals into a single control loop^7^.

VGICs are mechanosensitive^24, 58–62^, but the mechanisms of their mechanosensitivity remain poorly understood because of intrinsic structural and functional limitations. We used the bacterial voltage-gated sodium channel NaChBac as a model because it shares crucial structural and functional elements^25, 26^ with voltage-gated sodium channels (Na_V_s). We found that NaChBac^26^ is mechanosensitive, and impressively, the mechanosensitive responses of NaChBac closely resembled those of Na_V_1.5 (Figure 1), with forces increasing the peak currents and accelerating the kinetics. These were consistent with previous studies using macroscopic currents to examine mechanosensitivity in eukaryotic Na_V_s^10, 17^ and other VGICs^11, 63, 64^, which further strengthens NaChBac as a model to study eukaryotic VGICs. In response to physiological levels of mechanical stimuli traditionally used to stimulate mechanogated ion channel^65^, NaChBac populations, and single channels substantially increased their activity in a voltage-dependent manner (Figures 1 and 2). Force produced asymmetric rises in peak voltage-gated currents as the membrane potential depolarized to activate the channels. It is important to note that forces did not directly open NaChBac without voltage stimuli (Figure 1 and Figure 2), suggesting that mechanical force does not create new conformational states but rather impacts a single transition along the gating pathway. While whole-cell experiments proved informative, single-channel studies were required to test our hypotheses directly.

We removed NaChBac inactivation (NaChBac T220A)^27, 28^, which allowed us to zoom in on the mechanosensitivity of voltage-dependent activation. Using NaChBac T220A along with technical modifications and paired-stimulus configuration, which controlled for the known resting elevated mechanical tension in patch bilayers^34, 66^, we were able to resolve sub-pA NaChBac events with mechanical stimulation (Figures 2-5). Physiologically relevant patch suction modified NaChBac voltage-gating, reversibly increasing NaChBac voltage-dependent open probability (P_O_) in a dose-dependent fashion. This effect was indeed state-dependent, suggesting that applied forces have a state-specific effect on the Na_V_ channel, where the added mechanical energy appears to modify the energy landscape of gating but does not overcome voltage-gating^46, 67^.

To explain NaChBac mechanosensitivity, we propose the “mechanosensitive opening” (MSO) model, rather than “mechanosensitive activation (MSA), which features NaChBac pore opening as one strongly mechanosensitive transition (Figure 4). It is remarkable considering the MSO model’s simplicity that it could fit both whole-cell and single-channel data. The critical model discriminator was the force-induced change in the whole-cell voltage-dependent activation curve: increased maximum response and slope with an unchanged foot. We discriminated the two models by voltage-induced changes in the single-channel pressure-dependent activation curve. Finally, the MSO model explained the pressure-dependent changes in the pore opening. As observed in single-channel data at maximally activating voltages, suction shortened closed state lifetimes, suggesting that pressure destabilizes the closed state and ruling out non-specific pressure effects. While the structures responsible for voltage and force sensitivity may be distinct and function independently, from the kinetic mechanism standpoint, voltage and force sensitivities are state-dependent and intertwined: voltage acts on states C_1_ through C_5_, whereas tension acts on states C_5_ and O_6_. Consequently, channels must first activate by voltage before responding to tension. While simplified, this model captures the essence of the VGIC function and can apply to both prokaryotic and eukaryotic sodium channels.

Comparing the closed and open bacterial Na_V_ crystal structures shows the most extensive area changes are in the intracellular gate during the transition from closed to open^48, 52^. The bottom halves of S6 form the intracellular gate, working like hinges on a door latched by non-covalent interactions. Functional and modeling studies support the *swinging door* model: targeting S6 residues around the pore’s hinge impedes gating^50, 51, 68^, and pore opening led to a physical expansion of the inner leaflet, suggesting a palpable area expansion^69^. Consistent with these studies, electrophysiology and modeling show that S6 in the pore stores mechanical energy of gating^46, 70^. Therefore, determining a mechanosensitive site via mutagenesis to the voltage-sensor or pore domain offers conclusive evidence for the MSO model. We targeted both sites separately to differentiate between the effects of force on voltage sensors from those on the pore. The S4 positively charged residues that sense voltage are stabilized in the resting state within the lipid bilayer by counterbalancing acidic (negatively charged) residues^54^. By mutating one of these acidic residues (D93), we left-shifted the voltage-dependence of activation but otherwise did not change mechanosensitivity, confirming that voltage sensors do not significantly contribute to mechanosensitivity (Figure 5). Our functional data suggested that S6, forming a highly conserved component of the intracellular gate, might influence NaChBac mechanosensitivity. After testing several dead mutants, we found and mutated a conserved hydrophobic residue I228 in the S6 lining the channel pore. While I228G did not appreciably affect voltage-gating, it eliminated response to pressure, demonstrating that the pore is critical for mechanosensitivity (Figure 5). Thus, these results agree with structural and functional data showing significant in-plane area expansion during channel gating, support the *swinging door* model of VGIC pore gating, and suggest that force and voltage collaborate to gate NaChBac.

Since broad structural aspects of the intracellular gate appear conserved across VGICs from prokaryotes to eukaryotes^71, 72^, we surmise that VGIC mechanosensitivity may be generalizable. If mechanosensitivity were deleterious, it would likely not have reached the level of prevalence it has; nature would have selected for a different gating mechanism without cross-section expansion. Nevertheless, it is a ubiquitous property observed across many families of VGICs^24, 61^ and across each phylum, including unicellular to complex multicellular organisms. Perhaps archaic prokaryotic ion channels and sodium channels overall have developed maintained mechanosensitivity as their earliest *sense*^73^, and sodium channels maintained it under selective pressure.

How does membrane tension reach the NaChBac pore? In the *force-from-lipid* model, bilayers transduce mechanical energy directly into channel gating^43, 74, 75^. For the tensed bilayer to perform work (F⋅d) on the channel, conformational transitions leading to the open state must associate with in-plane area expansion during opening and contraction during closing^45^. Bilayers self-assemble to minimize contact between lipid tails and water molecules. However, despite minimization of free energy in assembled bilayers, the physical and energetic differences between phospholipid headgroups and lipid tails produce substantial intrinsic lateral forces^76^ reaching 1,000 atm^77^. These lateral forces act upon the protein-lipid interface of ion channels^65, 78^ and have non-homogeneous effects on resident proteins through the bilayer thickness: the hydrophobic lipid core applies compression while phospholipid head groups apply tension (Figure 6). Specialized mechano-gated ion channels are logical candidates to take advantage of this physical arrangement, and indeed they leverage forces developed at the protein-lipid interface for their *force-from-lipid* gating^43, 65, 78, 79^. For VGICs, both voltage sensors^80^ and pore-forming structures are bathed in phospholipids^81^. Therefore, it is reasonable to conclude that lipids could contribute to force sensing^11, 46^, given that lipids are crucial for voltage-dependent gating^80, 82^ and pore opening^10, 46, 74, 81^, and lipid permeable compounds frequently alter VGIC mechanosensitivity^21, 83^. Further work is required to determine the effects of lipid-protein interactions on VGIC mechanosensitivity.

**Figure 6.**
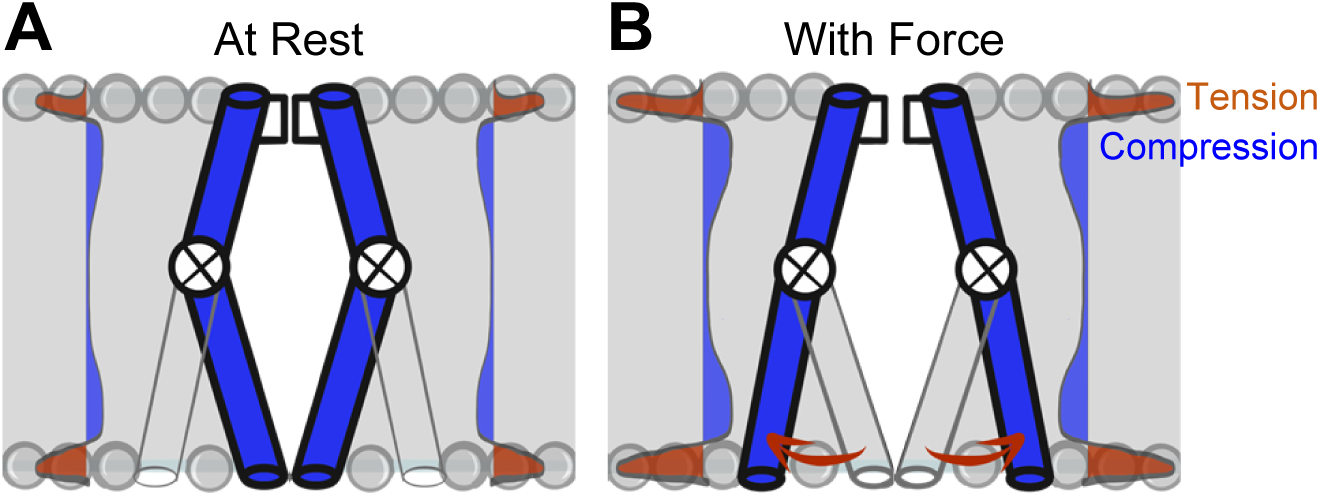
Model of voltage-gated ion channel mechanosensitivity. **(A)** VGIC pore is embedded in the lipid bilayer, which has an intrinsic distribution of mechanical forces even with no tension added to the system. **(B)** Mechanical stress applied to the bilayer alters the profile of bilayer forces, which destabilizes the intracellular gate and leads to intracellular pore expansion.

VGIC’s P_O_-dependent mechanosensitivity has important physiologic implications, allowing Na_V_ channels to serve as voltage-sensitive mechanosensors. Force can adjust the voltage set point for Na_V_ channel activation and affect action potential upstroke, regulating excitability^5, 6^.

Meanwhile, mechanosensitivity in voltage-gated potassium (K_V_) channels^11^ may serve as a mechanical brake on neuronal hyperexcitability in a voltage-sensitive fashion^7^. Beyond roles for VGIC mechanosensitivity in physiology, studies have uncovered patient VGIC mutations with functional disruptions in mechanosensitivity associated with diseases such as long-QT syndrome^18^ and irritable bowel syndrome (IBS)^20, 84^.

VGIC mechanosensitivity could be pharmacologically targeted in mechano-pathologies. Although specific VGIC mechanosensing inhibitors remain undeveloped, recent studies show that some amphipathic compounds Na_V_ channels are effective blockers of Na_V_ mechanosensitivity, separate from their local anesthetic mechanism^21, 22, 83^. Interestingly, the compounds’ amphipathic nature is critical for function^21, 83^, implying the channel pore’s lipid-protein interface is crucial for VGIC mechanosensitivity and suggesting the intracellular gate’s interaction with lipids may provide a novel pharmacologic target.

To summarize, we show here that the prokaryotic VGIC NaChBac is intrinsically mechanosensitive, and its mechanosensitivity depends on the channel pore intracellular gate. These results offer opportunities for future studies to determine roles for Na_V_ channel mechanosensitivity in physiology and pathophysiology and target Na_V_ mechanosensitivity in disease.

## MATERIALS AND METHODS

### Cell culture

Human embryonic kidney cells (HEK293; American Type Culture Collection, Manassas, VA) were cultured in minimum essential medium (MEM, 11095-080) supplemented with 10% fetal bovine serum (FBS, 10082147) and 1% penicillin-streptomycin (15140-122, Life Technologies, Co., Grand Island, NY). Regular or Piezo1 knockout (P1KO) HEK293 cells (a kind gift from Dr. Ardem Patapoutian, Scripps Research Institute^30^) were transfected with DNA plasmids encoding wild-type Na_V_1.5 (variant H558/Q1077del) or wild-type or T220A NaChBac, along with GFP as a reporter, by Lipofectamine 3000 reagent (L3000-008) in OPTI-MEM medium (31985-070; Life Technologies, Co., Grand Island, NY). Transfected cells were incubated at 37 °C for 24 h (Na_V_1.5) or 32 °C for 24-48 h (WT or T220A NaChBac). Then, cells were lifted by trypsin and resuspended in NaCl Ringer’s extracellular solution (composition below) before electrophysiology.

Site-directed mutagenesis was performed in the T220A NaChBac background to introduce an additional mutation, I228G or D93A, by using the QuikChange Lightning Site-Directed Mutagenesis Kit (Agilent Technologies, Santa Clara, CA). Upon verification of construct integrity and successful mutagenesis by DNA sequencing, either plasmid was transfected into P1KO cells for electrophysiology (Table 3, Figure 5).

**Table 3.**
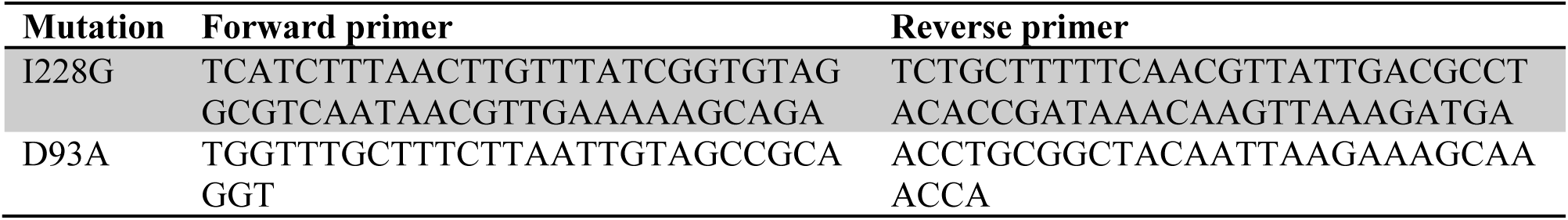
Primers for mutagenesis of I228G or D93A in the T220A NaChBac background.

### Electrophysiology

#### Pipette fabrication and data acquisition

Pipettes were pulled from KG-12 or 8250 glass (King Precision Glass, Claremont, CA) for whole-cell or cell-attached patches, respectively, on a P-97 puller (Sutter Instruments, Novato, CA) and coated with HIPEC R-6101 (Dow Corning, Midland, MI). Data were acquired with an Axopatch 200B amplifier, Digidata 1440A or 1550, and pClamp 10.6-11.2.1 software (Molecular Devices, Sunnyvale, CA).

#### Recording solutions

*For whole-cell electrophysiology of WT or T220A NaChBac*, the extracellular solution was NaCl Ringer’s, containing (in mM): 150 Na^+^, 5 K^+^, 2.5 Ca^2+^, 160 Cl^-^, 10 HEPES, 5.5 glucose, pH 7.35, 300 mmol/kg. The intracellular solution contained (in mM): 145 Cs^+^, 5 Na^+^, 5 Mg^2+^, 125 CH_3_SO_3_^-^, 35 Cl^-^, 10 HEPES, 2 EGTA, pH 7.0, 300 mmol/kg. *For whole-cell electrophysiology of Na_V_1.5* and *cell-attached patch clamp of T220A NaChBac*, the bath (extracellular) solution contained (in mM): 135 Cs^+^, 15 Na^+^, 5 K^+^, 2.5 Ca^2+^, 160 Cl^-^, 10 HEPES, 5.5 glucose, pH 7.35, 300 mmol/kg. The pipette solution for cell-attached patches was NaCl Ringer’s, supplemented with 0.03 mM Gd^3+^ to inhibit leak currents.

#### Whole-cell voltage clamp

Whole cell Na^+^ currents from HEK293 cells heterologously expressing Na_V_1.5 (variant H558/Q1077del) or WT or T220A NaChBac were recorded with a two-pulse protocol that tests channel activation during the first step and channel availability (steady-state inactivation) during the second step. Cells expressing Na_V_1.5 were pulsed every 1 s from the -130-mV holding potential through -10 mV in 5 mV intervals during step 1, then immediately pulsed to -40 mV for 50 ms during step 2. Na_V_1.5 data were sampled at 20 kHz and filtered at 5 kHz. Cells expressing NaChBac were pulsed every 4.75 s from the -120-mV holding potential through 0 mV in 10 mV intervals during step 1, then immediately pulsed to 0 mV for 50 ms (WT) or -50 mV for 400 ms (T220A) during step 2 (Figure 1 Suppl. A). NaChBac data were sampled at 2 kHz and filtered at 1kHz.

#### Cell-attached patch-clamp

P1KO cells heterologously expressing T220A NaChBac channels were held at -120 mV. To obtain single-channel events, we recorded thousands of sweeps in response to a voltage ladder protocol containing five 400 ms-long steps, from -100 mV to -20 mV in 20 mV increments, with a 3 s inter-sweep interval. Each voltage step was divided in two 200 ms-long pressure steps, from 0 mmHg to -10, -30, or -50 mmHg. Because the D93A mutant had open and closed times approximately 2-5 times longer than T220A, D93A experiments were performed with 4 s-long voltage steps and 2 s-long pressure steps. To test reversibility following pressure, the duration of each of the five voltage steps was 1 s with a 7.5 s inter-sweep interval, and pressure was applied for 500 ms (Figure 3 Suppl. A). Capacitance and passive currents were subtracted with a 1-sweep blank record, averaged from several to dozens of traces from the same or a subsequent recording in which no channel openings were observed^85^.

#### Mechanical stimulation

Mechanical stimuli were applied by shear stress to the entire cell, and by pressure clamp to membrane patches, as previously described^21, 22^. For whole-cell electrophysiology, shear stress was applied as the flow of extracellular solution through the 700- µL elliptical bath chamber, for 60-90 s at 10 mL/min^20, 22^. For cell-attached patch-clamp experiments, a negative pressure of -10 or -30 mmHg was applied by high-speed pressure clamp (HSPC-1, ALA Scientific Instruments, Farmingdale, NY)^31^. The single-channel data were sampled at 20 kHz and low-pass filtered on-line at 5 kHz but for analysis were further filtered at 0.5 kHz, due to a bandwidth limitation imposed by the HSPC (Figure 2 Suppl. G). The pressure clamp was set to +10 mmHg before the pipette entered the bath, then it was stepped to 0 mmHg after the tip contacted the cell membrane. Initial pipette resistance was 1-2 MΩ, and seal resistance was >10 GΩ.

### Data analysis

Data were analyzed in pClamp version 10.6 or 11.0.3 (Molecular Devices, Sunnyvale, CA), Excel 2010 (Microsoft, Redmond, WA), and Sigmaplot 12.5 (Systat Software, San Jose, CA). To estimate whole-cell conductance and voltage-dependent activation, the peak current evoked by voltage step 1 in the protocol described above was fit with a Boltzmann equation, *I_V_* = (*V* – *E_rev_*) x *G_Max_*/(1 + e^((*v*-v1/2*a*)/*δVa*)^), where *I_V_* is the peak current (pA/pF) at the test voltage

*V* (mV), *E_Rev_* is the reversal potential (mV), *G_Max_* is maximum conductance (nS), *V_1/2a_* is the half- activation voltage (mV), and *δV_a_* is the voltage sensitivity of activation (mV). To estimate voltage- dependent inactivation, the peak current *I_V_* evoked by voltage step 2 in the protocol was first normalized as a percentage to its maximum across all sweeps and then was fit with a Boltzmann equation, *I_V_* = 1/(1 + e^((*v_1_*-v_1/2_*i*)/*δV_i_*)^), where *V_1/2i_* is the half-inactivation voltage and *δV_i_* is the voltage sensitivity of inactivation. For kinetic analysis, whole-cell currents were fit to an exponential equation, *I_t_* = A_1_ × e^-t/^*^τa^* + *A*_2_ × e^-t/^*^τi^* + C, where *τ_a_* and *τ_i_* are activation and inactivation time constants (ms), respectively, and *A_1_*, *A_2_*, and *C* are constants.

To characterize single-channel conductance properties, all-point histograms of T220A NaChBac single-channel activity were fit with a sum of two Gaussian functions, 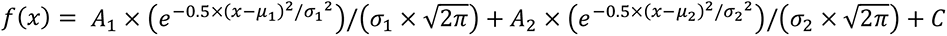, where *x* is current (pA), *µ* and *σ* represent the mean and standard deviation of the closed and open state current (pA), *A_1_* and *A_2_* are the weights of the closed and open state Gaussian components, respectively, and *C* is baseline current. Open probability was calculated as *P_O_=A_2_/(A_2_+A_1_)*. The response to pressure, P_O_(x)–P_O_(0), where x stands for -10 or -30 mmHg, was obtained as the difference in P_O_ values within the same trace. The single-channel closed and open times were calculated in pClamp 11.1 from idealized single-channel data. Data are expressed as means ± standard error (SEM). Change from shear stress or pressure was considered statistically significant when *P*<0.05 for mechano- stimulus vs. control, as determined by a 2-way ANOVA with Dunnett’s post-test.

#### Single-channel data analysis and simulations

The analysis and simulations were done with the QuB program, the MLab edition (http://milesculabs.org/QuB.html). QuB was used to digitally low-pass filter the data at 0.5 kHz to eliminate a periodic artifact induced by the pressure clamp system (Figure 2 Suppl. G) and to extract (“idealize”) the signal from the noisy data. QuB was further used to simulate the behavior of the tested NaChBac model and to calculate its properties: the voltage-activation curve at different pressures, the pressure-activation curve at different voltages, and the probability density function for closed and open dwell times, and to extract rate constants from single channel data, using a first-order approximation to correct for missed events^40^.

#### Na_V_ channel model

To capture the basic properties of the NaChBac channel (homotetramer, inactivation removed), we used the simple linear kinetic scheme C_1_-C_2_-C_3_-C_4_-C_5_- O_6_. Each rate constant had the general expression *k* = *k*_0_ × exp(*k*_v_ × *V* + *k*_p_ × *P*), where *V* is membrane potential, *P* is patch pressure, *k*_0_ is a pre-exponential factor representing the value of the rate constant at zero voltage and pressure, and *k*_v_ and *k*_p_ are sensitivity factors for voltage and pressure, respectively. Lack of voltage or pressure dependence was encoded by setting *k*_v_ or *k*_p_ to zero. The rates along the activation pathway were in the expected 4:3:2:1 ratio (e.g., *k*_23_ = 2 × *k*_45_). The parameters of the model were tweaked by hand to match the macroscopic and single-channel data. First, we chose a set of *k*_0_ preexponential parameters for the C_5_-O_6_ transition, to match the observed P_O_ at saturating voltages (at -20 mV). Then, we adjusted the *k*_v_ exponential parameters that describe the voltage sensitivity of the C_1_ through C_5_ transitions, to match the normalized macroscopic activation curve under no-shear conditions. Next, we determined the statistical distribution (average and standard deviation) of the resting potential of the single-channel patched cells—to match the voltage-dependent P_O_ curve—which is voltage-shifted and shallower relative to the macroscopic activation curve. To generate a P_O_ curve that takes into account the variable and non-zero resting potential, the P_O_ value at each voltage point was obtained by numerically integrating over the Gaussian distribution describing the resting potential. Next, we adjusted the *k*_0_ preexponential parameters for the C_1_ through C_5_ transitions to approximately match the observed single-channel lifetimes. Finally, for the MSO model, we adjusted the *k*_p_ exponential parameters describing the pressure sensitivity of the C_5_ to C_6_ transition, to match the P_O_ curve under negative patch pressure. The same *k*_p_ values were also used for the MSA model.

## ACKNOWLEDGEMENTS

We would like to thank Drs. Simone Mazzaferro, Steven Sine, Paul DeCaen, Fred Sachs, Mirela Milescu, and Corrie DaCosta for their constructive suggestions, Denika Mueller for technical assistance, and Kristy Zodrow for administrative assistance. L.S.M. acknowledges the gracious support provided by Dr. Sergei Sukharev and the University of Maryland at College Park. NIH DK052766, DK123549, AT010875.

## AUTHOR CONTRIBUTIONS

<u>Peter R. Strege</u>: conceived and designed research, performed experiments, analyzed data, interpreted results of experiments, prepared figures, drafted manuscript, edited and revised manuscript, approved the final version of the manuscript

<u>Luke M. Cowan</u>: performed experiments, analyzed data, interpreted results of experiments, edited and revised manuscript, approved the final version of the manuscript

<u>Amelia Mazzone</u>: performed experiments, approved the final version of the manuscript

<u>Constanza Alcaino</u>: performed experiments, approved the final version of the manuscript

<u>Christopher A. Ahern</u>: conceived and designed research, edited and revised manuscript, approved the final version of the manuscript

<u>Lorin S. Milescu</u>: conceived and designed research, wrote analysis scripts, interpreted results of experiments, edited and revised manuscript, approved the final version of the manuscript

<u>Gianrico Farrugia</u>: conceived and designed research, interpreted results of experiments, edited and revised manuscript, approved the final version of the manuscript

<u>Arthur Beyder</u>: conceived and designed research, analyzed data, interpreted results of experiments, drafted manuscript, edited and revised manuscript, approved the final version of the manuscript

## SUPPLEMENTARY FIGURE LEGENDS

**Figure 1 Supplement.**
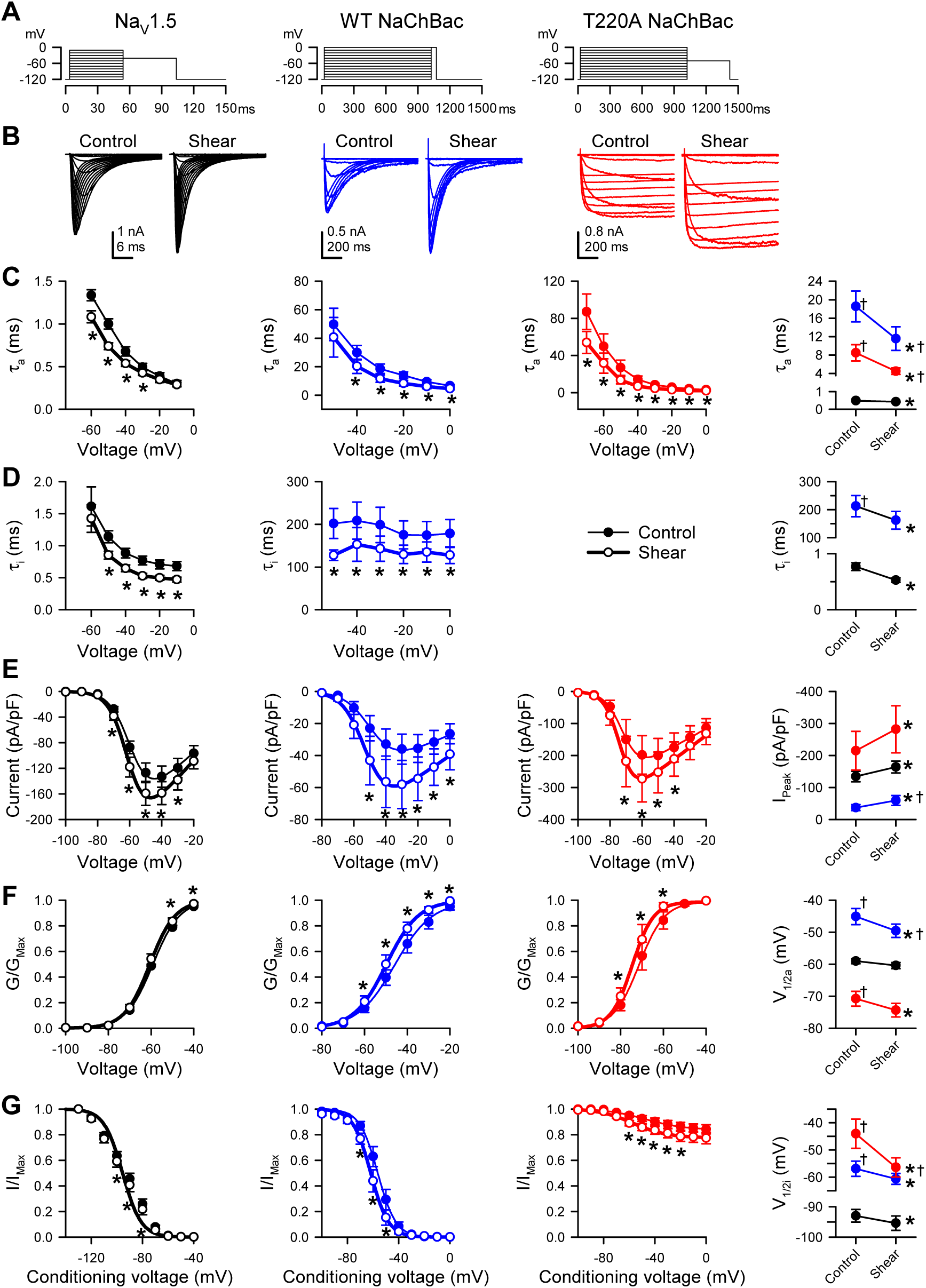
Shear stress increases peak Na^+^ current, hyperpolarizes the voltage- dependence, and accelerates the kinetics of eurkaryotic and prokaryotic Na_V_ channels in HEK293 cells. (A-B) Voltage protocols (A) elicited currents (B) from Na_V_1.5 and WT or T220A NaChBac channels transiently expressed in HEK293 cells. Currents were recorded before (*control*) or during (*shear*) flow of bath (extracellular) solution through the recording chamber at a rate of 10 mL/min. **(C-D)** Time constants of activation (C, τ_a_) or inactivation (D, τ_i_) versus step voltage, before (●) or during (○) shear stress. **(E)** Current density-voltage relationship of peak Na^+^ currents, before (●) or during (○) shear stress. **(F-G)** Steady-state voltage dependence of activation (F) and availability (G), recorded before (●) or during (○) shear stress. *Far right column*, mean parameters for the time constants of activation (C, τ_a_) or inactivation (D, τ_i_) at -30 mV, the maximum peak Na^+^ current (E, I_Peak_), the half-point of steady-state activation (F, V_1/2a_), and the half-point of steady-state availability (G, V_1/2i_), recorded from paired controls (Control) or with shear stress (Shear). Voltage clamp data were recorded from n = 7-10 cells each; *P<0.05 to control or †P<0.05 to Na_V_1.5 by two-way ANOVAs with Dunnett’s post-test.

**Figure 2 Supplement.**
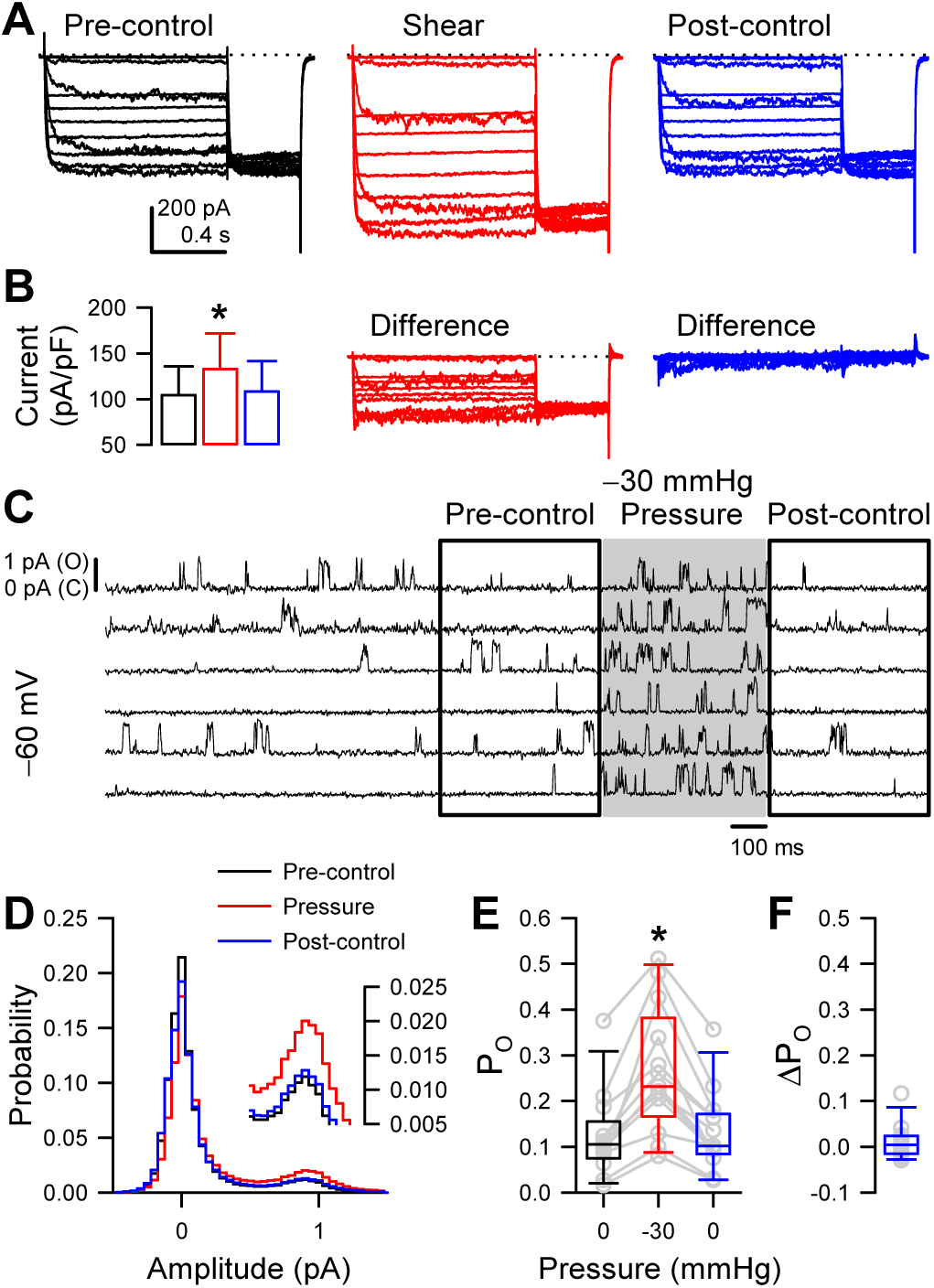
Endogenous channels in Piezo1-KO HEK (P1KO) cells are insensitive to pressure stimulus. **(A)** Single channel activity from an untransfected P1KO cell before or during (shaded area) application of pressure by high-speed pressure clamp (HSPC). **(B)** All-sample distribution curves generated from all traces recorded from the cell represented in (A), at +60 mV and with 0 (black) or -30 mmHg pressure stimulus (red). **(C)** Voltage- and pressure-clamp protocols to test the pressure sensitivity of single channel currents to -30 mmHg at voltage steps from -60 through +100 mV. **(D)** Single channel currents averaged from 60 sweeps of the protocol shown in (C)—a holding voltage of -100 mV to steps from -50 to +100 mV with 0 (control) or -30 mmHg (pressure) applied to the patch. **(E)** Difference current obtained by subtracting pressure from control currents in (D) (I_Difference_ = I_Control_ – I_Pressure_). **(F)** Current-voltage (I-V) relationship from control (black symbols), pressure (red), or difference (white) currents at the plateau, as shown in (D-E). (Inset) Enlargement of currents from -60 to 0 mV. **(G)** Noise spectrum averaged from 25 ten-second traces without (black) or with (red) the high-speed pressure clamp (HSPC) connected to the patch clamp headstage. Vertical gray lines indicate multiples of 60 Hz. Noise exclusive to HSPC ≈ 1.7 kHz.

**Figure 3 Supplement.**
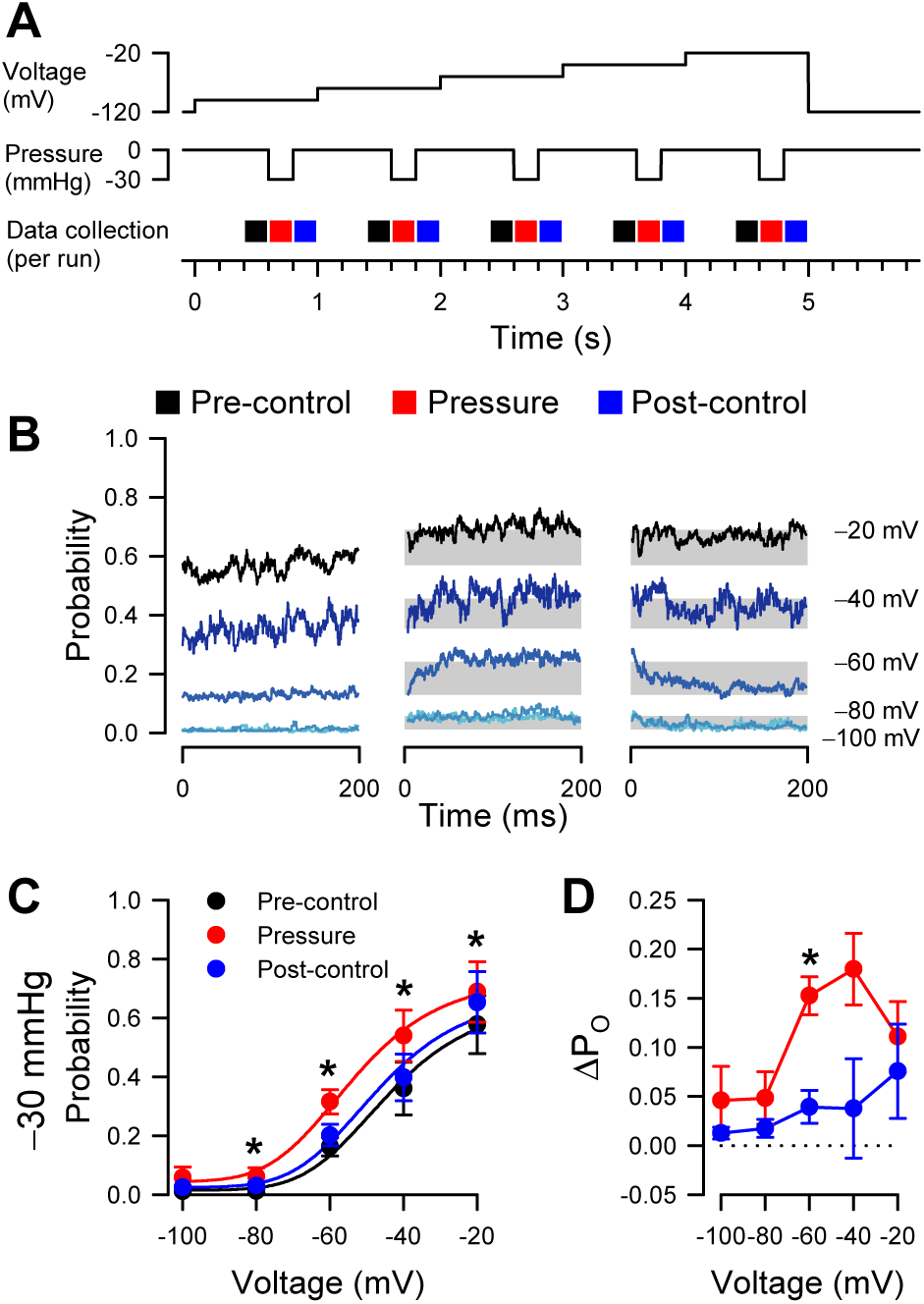
Effect of pressure on voltage dependent open probability. **(A)** Protocols to test reversibility of pressure-dependent increases in P_O_. **(B)** Current traces averaged from idealized single channel events in 4-17 cells at voltage steps from -100 to -20 mV, before (black), during (red), or after (blue) the pressure step to -30 mmHg. Shaded areas represent the difference in average P_O_ with pressure versus each pre-control baseline. **(C)** Single channel open probability versus voltage. **(D)** Differences in open probability (ΔP_O_), subtracting the open probability before pressure from either pressure (red) or post-control (blue).

**Figure 5 Supplement.**
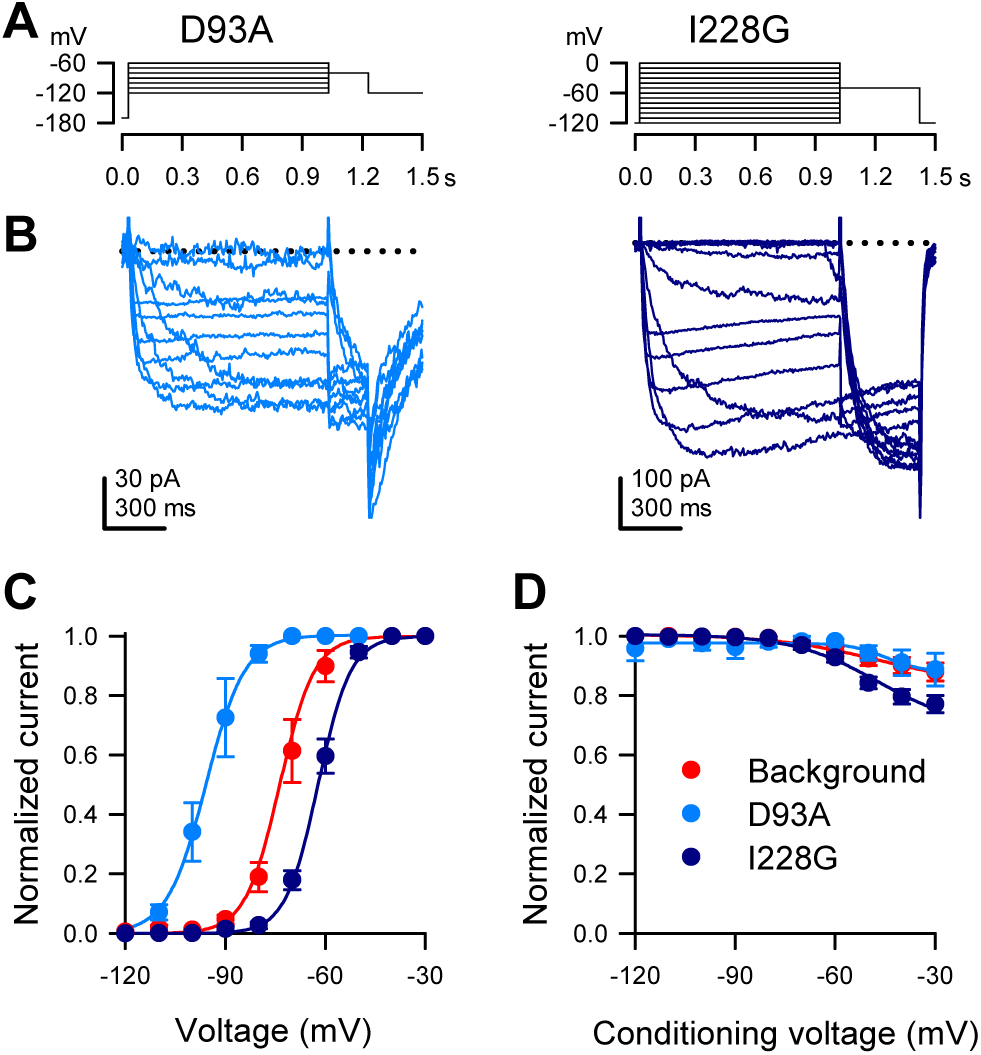
Whole cell voltage-dependent Na^+^ currents elicited from P1KO cells transfected with NaChBac mutants D93A or I228G in the T220A background. **(A)** Voltage stimulus protocol to elicit whole cell Na^+^ currents by holding the cell at -170 (D93A) or -120 mV (I228G), then stepping to a voltage ladder from -120 through -60 (D93A) or through 0 mV (I228G) for 1 s, then to a single voltage at -80 mV for 200 ms (D93A) or -50 mV for 400 ms (I228G). **(B)** Whole cell Na^+^ currents elicited by the voltage protocols shown in (A). (C) Steady-state activation curves versus the voltage of step 1 for the T220A background (red) or the mutants D93A (blue) or I228G (indigo). (C) Steady-state availability (inactivation) currents at step 2 versus the conditioning voltage of step 1 for background (red) or mutant D93A (blue) or I228G (indigo) channels (n = 8 (T220A), 3 (D93A), or 11 (I228G) transfected P1KO cells)

